## Supplementary material for "Circumventing the optical diffraction limit with customized speckles": Supplimentary


### 1 Experimental setup for photoconversion

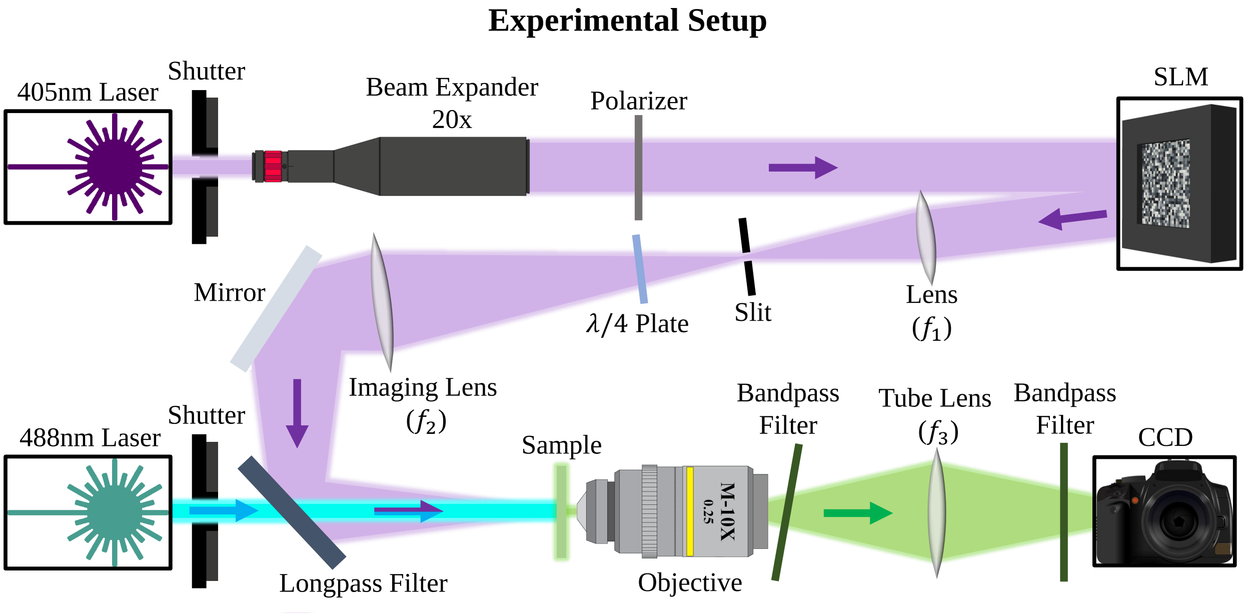


Figure 1: **Experimental setup.** The experimental setup used for nonlinear speckle-illumination microscopy is shown. We use a phase-only SLM to generate a customized speckle pattern to illuminate a fluorescent sample for photoconversion.

Figure 1 is a schematic of our experimental setup for photoconverting a fluorescent sample with customized speckle patterns and imaging the unconverted fluorescence. A CW laser operating at a wavelength of $\lambda = 405$ nm is used to photoconvert the mEos3.2 protein sample. The laser beam is expanded and linearly polarized before it is incident on a phase-only SLM (Meadowlark Optics). The pixels on the SLM can modulate the phase of the incident field between $0$ and $2\pi$: in increments of $2\pi/90$. Because a small portion of light reflected from the SLM is unmodulated, we write a binary phase diffraction grating on the SLM and use the light diffracted to the first order for photonconversion. In order to avoid cross-talk between the neighboring SLM pixels, $32 \times 32$ pixels are grouped to form one macropixel, and the binary diffraction grating is written within each macropixel with a period of 8 pixels. We use a square array of $32 \times 32$ macropixels in the central part of the phase modulating region of the SLM to shape the photoconverting laser light. The light modulated by our SLM is Fourier transformed by a lens with a focal length of $f_1 = 500$ mm and cropped in the Fourier plane with a slit to keep only the first-order diffraction. The complex field on the Fourier plane of the SLM is imaged onto the surface of the sample by a second lens, with a focal length of $f_2 = 500$ mm. Using a $\lambda/4$ plate, we convert the linearly polarized photoconverting beam into a circularly polarized beam before it is incident upon the sample. In this setup, the full width at half-maximum of a diffraction-limited focal spot is 17 µm.

After the photonconverion process, a second laser, operating at a wavelength of $\lambda = 488$ nm, uniformly illuminates the sample and excites the non-photoconverted mEos3.2 proteins. To collect the fluorescence, we use a $10\times$ objective of $\mathrm{NA} = 0.25$ and a tube lens with a focal length of $f_3 = 150$ mm. The 2D fluorescence image is recorded by a CCD camera (Allied Vision

Manta G-235B). The spatial resolution of the detection system is estimated to be 1.1 μm. The remaining excitation laser light, which is not absorbed by the sample, is subsequently removed by two Chroma ET 535/70 bandpass filters. One filter is placed after the objective lens and reflects the excitation beam off the optical axis of our system. The second one is placed directly in front of the camera.

### 2 Live sample demonstration

In the main text, we use delta speckle patterns to photoconvert a homogeneous film of purified protein to demonstrate the isometric and isotropic spatial resolution enhancement they allow. Here we show that our technique is compatible with living samples, which are typically inhomogeneous.

**Live yeast cell culture and preparation.** The *S. Pombe* strain *Leu1::Leu1+ pAct1 mEos3.2 nmt1Term ade6-M216 his3-Δ1 leu1-32 ura4-Δ18* is generated from NruI digested plasmid pJK148-pAct1-mEos3.2-nmt1Term through homologous recombination. Cytosolic mEos3.2 is expressed from the act1 promoter in the endogenous Leu1 locus. Cells are grown in exponential phase at 25 °C in YE5S-rich liquid medium in 50 mL flasks in the dark before switching to EMM5S-synthetic medium for 12-18 hours, to reduce the cellular autofluroescence background. Live cells are concentrated 10- to 20-fold by centrifugation at 3,000 rpm for 30 s and resuspended in EMM5S for imaging. Concentrated cells in 10 uL are mounted on a thin layer consisting of 35 μL 25 % gelatin (Sigma-Aldrich; G-2500) in EMM5S.

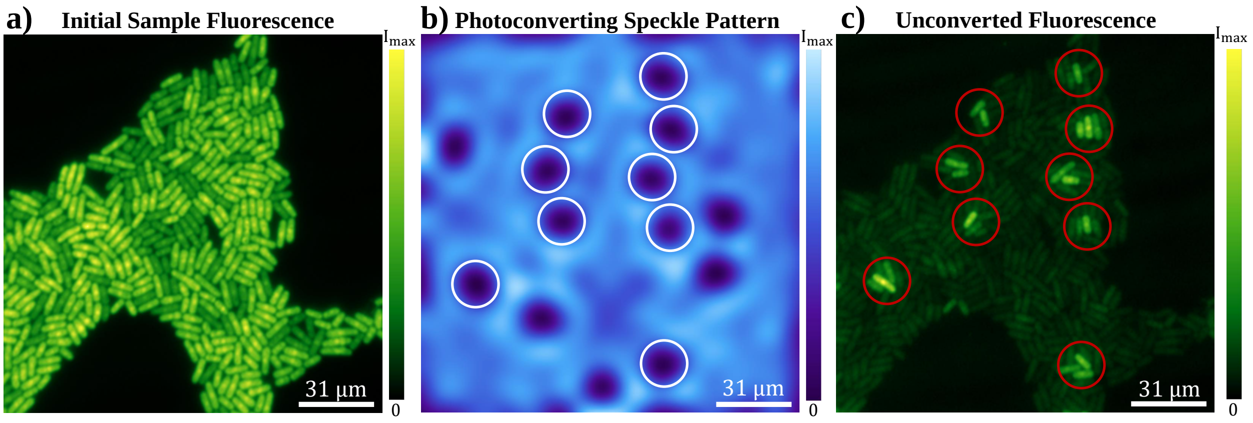


Figure 2: **Photoconversion of live yeast cells with a customized speckle pattern.** In **a**, we present an optical image of the fluorescent light emitted from a sample of live yeast cells before they are photoconverted by the delta speckle pattern shown in **b**. The white circles in **b** indicate the vortices which overlap with the yeast cells. An image of the fluorescent light emitted by the live-cell sample after photoconversion is shown in **c**. The red circles in **c** correspond to the white circles in **b**.

**Photonconverting live yeast cells with customized speckles.** We illuminate the collection of nonuniformly distributed yeast cells shown in Fig. 2**a** with the photoconverting speckle pattern presented in **b**. The optical vortices of the delta speckle pattern do not photoconvert the yeast cells in their vicinity. Consequently after photoconversion, multiple isolated groups of cells will emit fluorescence when illuminated by 488 nm light. In the fluorescence image taken after photoconversion, the fluorescent regions (marked by red circles in **c**) have a one-to-one correspondence to the optical vortices -in the delta speckle pattern- that overlap with the yeast cells (marked by white circles in **b**). Because the mEos3.2 protein is cytosolic, it is impossible to obtain fluorescence from a sub-cellular region, even if we shrink the dark region surrounding each vortex core using high-NA optics. Nevertheless, we are able to select individual groupings of live cells located near the vortices.

### 3 The joint PDF of the complex speckle fields

A speckle pattern is 'fully developed' if the joint PDF of its complex field is azimuthally invariant. This means that the phase distribution of the speckle pattern is uniform between $0$ and $2\pi$, and that the amplitude values of the field are independent of the phase values. This means that the amplitude and phase distributions of a fully-developed speckle pattern are statistically independent. In Fig. 3, we plot the complex field PDF calculated from 1,000 Rayleigh speckle patterns and 1,000 delta speckle patterns. While the field PDF obeyed by Rayleigh speckles is a circular Gaussian function, the delta speckles adhere to a circular non-Gaussian function. Because the field PDF is circular for delta speckles, they are fully developed like Rayleigh speckles.

### 4 Speckle decorrelation upon axial propagation

The rapid axial decorrelation of a fully-developed speckle pattern enables parallel 3D nonlinear patterned-illumination microscopy. The axial intensity correlation function is defined as

$$C_I(\Delta z) \equiv \frac{\langle \delta I(\mathbf{r}, z_0)\delta I(\mathbf{r}, z_0 + \Delta z)\rangle}{\sqrt{\langle[\delta I(\mathbf{r}, z_0)]^2\rangle}\sqrt{\langle[\delta I(\mathbf{r}, z_0 + \Delta z)]^2\rangle}} \tag{1}$$

where $z_0$ is the axial position of the focal plane, $\langle ... \rangle$ denotes averaging over transverse position $\mathbf{r}$, and $\delta I(\mathbf{r}, z) = I(\mathbf{r}, z) - \langle I(\mathbf{r}, z)\rangle$ is the intensity fluctuation around the mean. The FWHM of $C_I(\Delta z)$ defines the axial decorrelation length. For a Rayleigh speckle pattern, the axial decorrela-

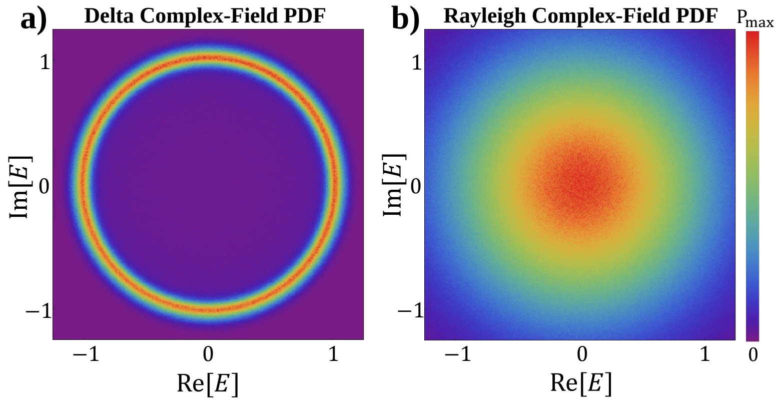


Figure 3: **Joint PDF of the complex speckle fields.** In **a**, we show the circular non-Gaussian joint PDF associated with delta speckles next to the circular Gaussian joint PDF of Rayleigh speckles in **b**. In both cases, the speckles are fully developed.

tion length $R_l$ is proportional to the Rayleigh range: which is determined by the axial diffraction limit $2\lambda/\mathrm{NA}^2$.

We numerically simulate the axial propagation of delta speckles and compare it with that of Rayleigh speckles. In Fig. 4, an example delta speckle pattern at $\Delta z = 0$ and $\Delta z = R_l/2$ is shown in **a** and **b**, while an example Rayleigh speckle pattern at $\Delta z = 0$ and $\Delta z = R_l/2$ is presented in **c** and **d**. A qualitative comparison between **a**, **b** and **c**, **d** illustrates that the delta speckles' transverse intensity-profile axially evolves much faster than the Rayleigh speckles' profile.

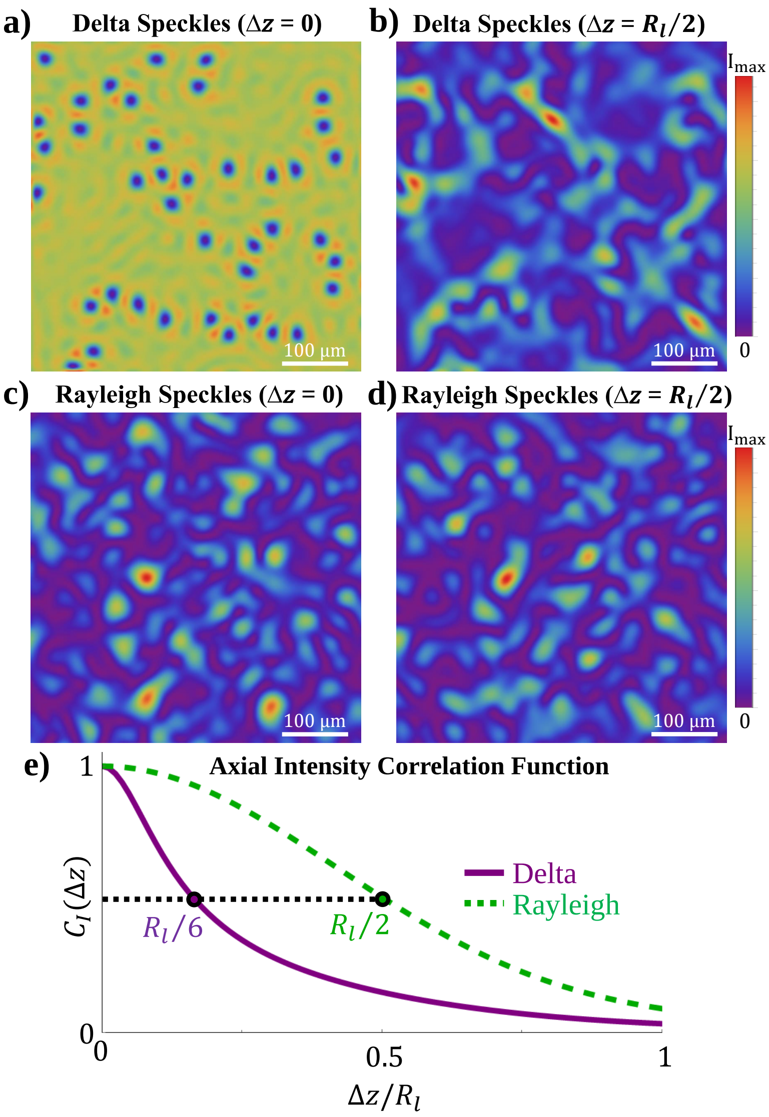


Figure 4: **Axial decorrelation of speckle patterns.** An example delta speckle pattern at $\Delta z = 0$ and $\Delta z = R_l/2$ is shown in **a** and **b**, while an example Rayleigh speckle pattern at $\Delta z = 0$ and $\Delta z = R_l/2$ is shown in **c** and **d**. In **e**, the axial intensity correlation function $C_I(\Delta z)$ for delta speckles (purple line) and Rayleigh speckles (green dashed line) is shown. We ensemble average over the propagation of 100 speckle patterns to create the curves in **e**.

We plot the axial correlation function of the delta speckles (purple solid line) and the Rayleigh speckles in **e**. The axial decorrelation length of the delta speckles (FWHM of axial intensity correlation function) is $R_l/3$, which is three times shorter than that of the Rayleigh speckles. As mentioned in the main text, the vortices in a delta speckle pattern deviate the most from the mean intensity value, thus they dictate the intensity correlation function. The accelerated axial decorrelation is partially attributed to the rapid transverse motion of optical vortices upon axial propagation. This characteristic of the delta speckles can enhance the axial resolution in 3D imaging.

In Fig. 5 we show the effect the transverse motion of optical vortices in delta speckles - upon axial propagation over one $R_l$- has on the fluorescent spots in a photoconverted sample. An example delta speckle pattern, $I(x, y, \Delta z = 0)$, is shown in **a** along with two vertical cross-sections, $I(x = x_0, y, \Delta z)$ in **b** and $I(x, y = y_0, \Delta z)$ in **c**. The axial cross-section distributions show the speckle's intensity, along a line in $x$ or $y$ axis, as a function of axial propagation from $\Delta z = 0$ to $\Delta z = R_l$. Using Eq. 2 from the main text, we simulate the corresponding unconverted protein density, $\rho(\mathbf{r}, t)$, in a sample after it is photoconverted with the speckle pattern shown in **a-c**. In **d-f** we show the unconverted protein densities after the sample is photoconverted for $t = 10/q$. Not only do the fluorescent spots located at $\mathbf{r} = \alpha, \beta, \gamma$ move transversely and leave the axial cross-section in **e** & **f** before propagating to $\Delta z = R_l$, but also, other fluorescent spots enter and exit the axial cross-sections: demonstrating that delta speckles can be used to obtain 3D super-resolution images.

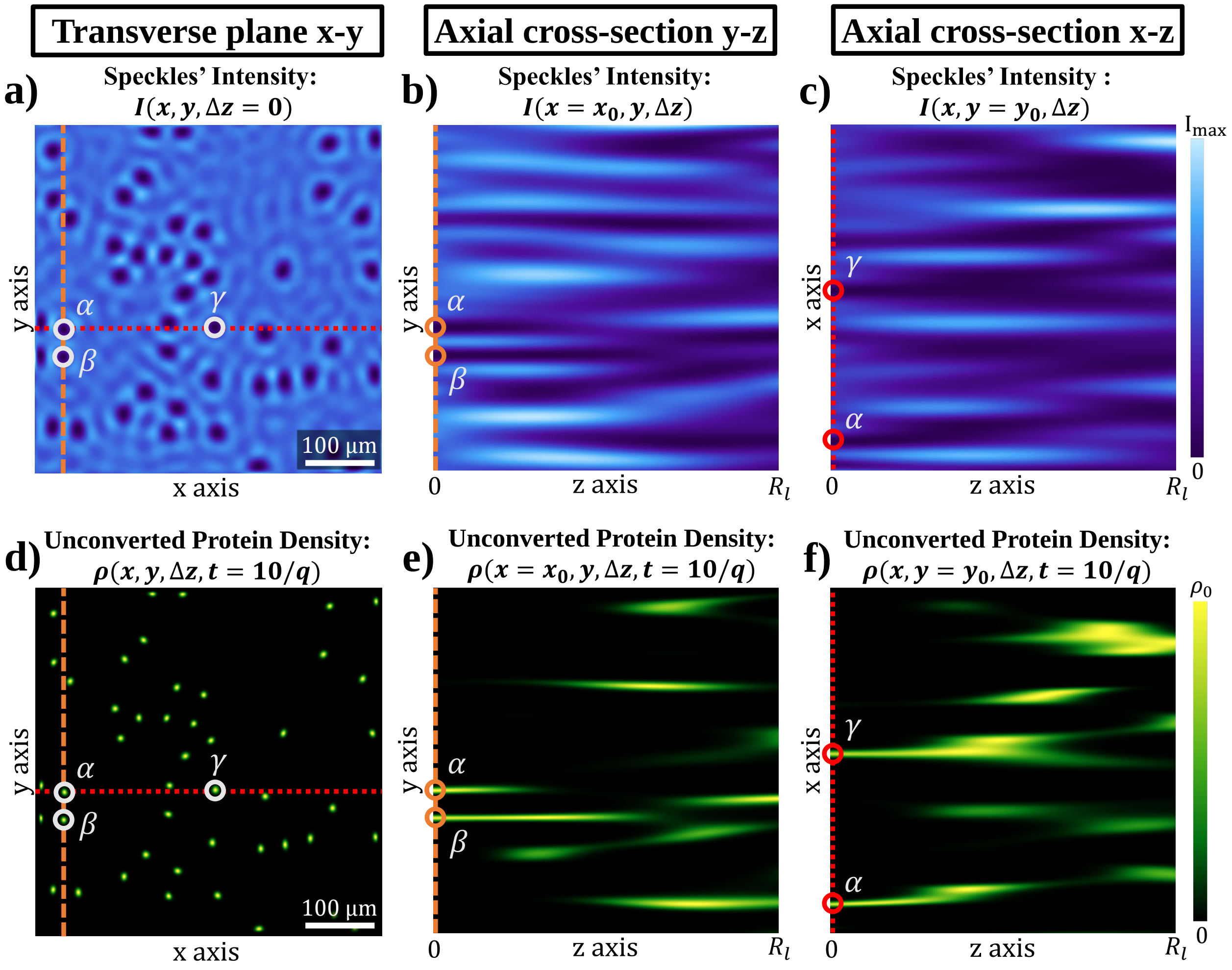


Figure 5: **Three-dimensional photoconversion with delta speckles.** The calculated spatial distribution of a delta speckle pattern's intensity on a transverse plane at $z = 0$, $I(x, y, \Delta z = 0)$, is shown in **a** along with the intensity distribution over two axial cross-sections, $I(x = x_0, y, \Delta z)$ in **b** and $I(x, y = y_0, \Delta z)$ in **c**. A photoconversion simulation using the speckles in **a**-**c** gives the corresponding distribution of the unconverted protein density in a uniform sample, $\rho(\mathbf{r}, t)$, in **d**-**f**.
